# ConvCGP: A Convolutional Neural Network to Predict Genotypic Values of Rice Traits from Compressed Genome-Wide Polymorphisms

**DOI:** 10.1101/2024.11.20.624609

**Authors:** Tanzila Islam, Chyon Hae Kim, Hiroyuki Shimono, Akio Kimura, Hiroyoshi Iwata

## Abstract

The growing size of genome-wide polymorphism data in animal and plant breeding has raised concerns regarding computational load and time, particularly when predicting genotypic values for target traits using genomic prediction. Although several deep learning and conventional methods, including dimensionality reduction techniques, such as principal component analysis (PCA) and autoencoders, have been proposed to address these challenges by selecting subsets of polymorphisms or compressing high-dimensional data for predictive analysis. However, these methods are often computationally intensive and time-consuming. A major challenge in applying high-dimensional genomic data directly to deep-learning models is the substantial computational cost and time required for hyperparameter tuning and model training. To address these limitations, we propose a novel deep learning approach that combines convolutional neural networks (CNNs) to predict the genotypic data of target traits with autoencoders to compress high-dimensional genome-wide polymorphism data. We tested this framework on high-dimensional rice datasets, focusing on agronomic trait prediction. By combining CNNs with autoencoders, our framework outperformed other machine-learning methods and recently proposed compression methods, demonstrating its potential to efficiently address the computational challenges associated with high-dimensional genomic data.

## 1 Introduction

Accurately estimating and predicting genotypic values underlying target traits is a central and ongoing challenge in plant and animal breeding (Lourenço et al., 2024). Genomic selection (GS) (Meuwissen et al., 2001), which uses genome-wide marker information to predict genotypic values and select superior genotypes (individuals or lines), has been proposed to significantly enhance breeding speed and efficiency (Hickey et al., 2017). Initially, GS relied on genotyping arrays, but with the advent of next-generation sequencing (NGS) (Marudamuthu et al., 2023), the resulting data have become increasingly high-dimensional. Managing and leveraging high-dimensional data for tasks, such as building prediction models, remains a significant challenge. As the number of samples (plants and animals) used in GS continues to increase, addressing these challenges will become even more critical.

Various strategies have been proposed for genomic prediction, regardless of the type of genomic data. Among these, deep learning (DL) methods have attracted significant attention in computational biology (Angermueller et al., 2016; Montesinos-López et al., 2021). DL has been applied in genome-wide association studies to identify SNP interactions (Uppu et al., 2016) and classify genomic variants (Liang et al., 2016). For example, DeepGS, an ensemble of convolutional neural networks (CNN) (Krizhevsky et al., 2017) and rrBLUP, has been used to predict genotypic values from imputed SNPs (Ma et al., 2018), whereas simple dense neural networks (DNN) have been utilized for genotype-by-sequencing (GBS) data (Montesinos-López et al., 2018). Abdollahi-Arpanahi *et al*. compared the predictive performances of two DL methods (multilayer perceptron [MLP] and CNN), two ensemble methods (random forest [RF] and gradient boosting), and two parametric methods (genomic best linear unbiased prediction [GBLUP] and BayesB) using real and simulated datasets (Abdollahi-Arpanahi et al., 2020). They found that DL methods only slightly outperformed parametric methods on large datasets. In these genotypic value prediction tasks, CNNs can capture spatial information from raw sequencing reads or genomic variants without the need for feature engineering. However, these DL methods face challenges in fully addressing the complexities of high-dimensional data analysis.

Recently, Nazzicari et al. proposed a dimensionality reduction method that uses kinship matrices to compactly represent SNP genotype data. Multiple kinship matrices were stacked and processed using a 2D-convolutional neural network for genetic value prediction (Nazzicari and Biscarini, 2022). Using a similar approach, Kick et al. applied principal component analysis (PCA) for dimensionality reduction and fed the data into a multimodal DL model. They found that the performance of multimodal DL was comparable to that of GBLUP (Kick et al., 2023). Although stacking kinship matrices and PCs helps reduce the dimensionality of the dataset, these methods may lose some fine-grained genetic information during compression, leading to inconsistent prediction performance when applied to compressed data. In our previous study, DeepCGP addressed the issue of high-dimensional data by applying autoencoders and random forest regression for compression-based genomic predictions (Breiman, 2001; Liaw and Wiener, 2002). DeepCGP successfully reduces the data dimensions while retaining moderate prediction performance using the random forest regression method (Islam et al., 2023).

In this study, we developed a deep-learning-based method called Compression-based Genomic Prediction using a Convolutional Neural Network (ConvCGP), which utilizes compressed information generated by autoencoders to predict the genotypic values of a target trait. The key contributions of this work are as follows. First, we demonstrate how high-dimensional genome-wide polymorphism data can be effectively compressed using autoencoders. Second, we demonstrate how compressed data can be applied to predict genotypic values using a CNN. To assess the performance of ConvCGP, we employed two rice genome datasets: C7AIR, with 7,098 single-nucleotide polymorphisms (SNPs), and HDRA, with 700,000 SNPs. Our results showed that ConvCGP accurately predicted genotypic values using compressed genome-wide polymorphism data, achieving a performance comparable to that of predictions made using the original, uncompressed data. Furthermore, we evaluated the compression-based prediction against other methods, highlighting the potential of deep-learning regression techniques for genomic prediction.

## 2 Materials and Methods

### 2.1 Datasets and Data Pre-processing

We used two datasets of varying sizes to illustrate the broad applicability of our models. We assessed the performance of the models by measuring how accurately the compressed genome-wide polymorphism data could predict the genotypic values of a target trait using a deep learning-based regression method. This study aimed to achieve a prediction performance comparable to that of models using original data while also surpassing other state-of-the-art approaches.

#### C7AIR

The first dataset, the Cornell-IR LD Rice Array (C7AIR) (Morales et al., 2020), is a second-generation SNP array containing 189 rice accessions for 7,098 markers from the Rice Diversity project. These accessions carried the estimated genotypic values for plant height.

#### HDRA

The second dataset, High-Density Rice Array (HDRA) (McCouch et al., 2016), consisted of 1,568 diverse inbred rice varieties with 700,000 SNPs. The genotypic averages of 34 traits were estimated for 388 lines (Zhao et al., 2011), although some records were missing. We excluded 29 genotypes with ten or more missing data points, leaving 359 lines across 18 traits for analysis. The genotype data were formatted as a bed matrix derived from the VCF format, where each entry was scored as 0, 1, or 2 to represent the genotype at each SNP: 0 for the homozygous reference allele, 2 for the homozygous alternate allele, and 1 for a heterozygous state (combinations of different nucleotides, such as AT, AC, AG, TC, TG, and CG). Given that all accessions were inbred lines, expected to be homozygous at most loci, we treated 1 as a missing value and converted 0 and 2 into categorical values (A, C, G, and T). The converted data were saved in the CSV format. We used the ‘gaston’ package (Perdry and Dandine-Roulland, 2018) in R for this conversion.

Categorical values (A, C, G, and T) were preprocessed in both datasets by applying one-hot encoding. Each genome was encoded into one-hot encoding using a 4-bit coding scheme, where **x**∈ R^d^×4, with d represented the length of the genome sequence. The nucleotides “A,” “C,” “G,” and “T” were encoded by “1000”, “0100”, “0010”, and “0001”, respectively. The C7AIR and HDRA datasets contained approximately 13% and 10% missing genotypes, respectively. Therefore, we encoded the missing values “N” by “0000.”

After one-hot encoding of the raw data, the dimensions of the C7AIR and HDRA data were 189 × 28392 and 1568 × 2800000, respectively. Owing to the high dimensionality of the input data, we applied a data splitting technique to reduce the computational time. Using the hsplit function of NumPy, the one-hot encoded arrays were split horizontally (along axis 1) into smaller chunks. For C7AIR, each split resulted in data with a size of 189 × 28, whereas for HDRA, each split was 1,568 × 28, creating input layers of 28 neurons for each network. Consequently, 1,014 autoencoder networks were used for the C7AIR dataset and 100,000 autoencoder networks for the HDRA dataset, respectively.

### 2.2 Compression Model using Deep Autoencoder

To compress genome-wide polymorphism data, we utilized a deep autoencoder (Kramer, 1991; Goodfellow et al., 2016) (Figure 1) based on two symmetrical deep belief networks with multiple hidden layers: (i) an encoder network ***h*(*x***_*i*_), where ***x***_*i*_ ∈ *R*^*d*^, which first encodes an input ***x***_*i*_ into a hidden representation ***h*(*x***_*i*_)^( *l*+1)^as shown Equation 1, and (ii) a decoder network ***x***′_*i*_, which maps the hidden representation ***h*(*x*_i_)**^( *l*+1)^ back into a reconstruction ***x***′ ^( *l*)^ as defined in Equation 2:

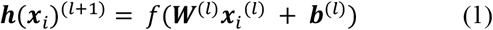

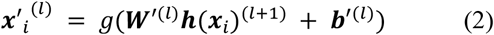

Where *f* is the encoding activation function, ***W***^( *l*)^is the encoding weight matrix, ***b***^( *l*)^is the encoding bias vector, *g* is the decoding activation function, ***W***′^( *l*)^is the decoding matrix, and ***b***′^( *l*)^ is the decoding bias vector from the *l*-th input layer to the ( *l* + 1)-th hidden layer.

**Figure 1.**
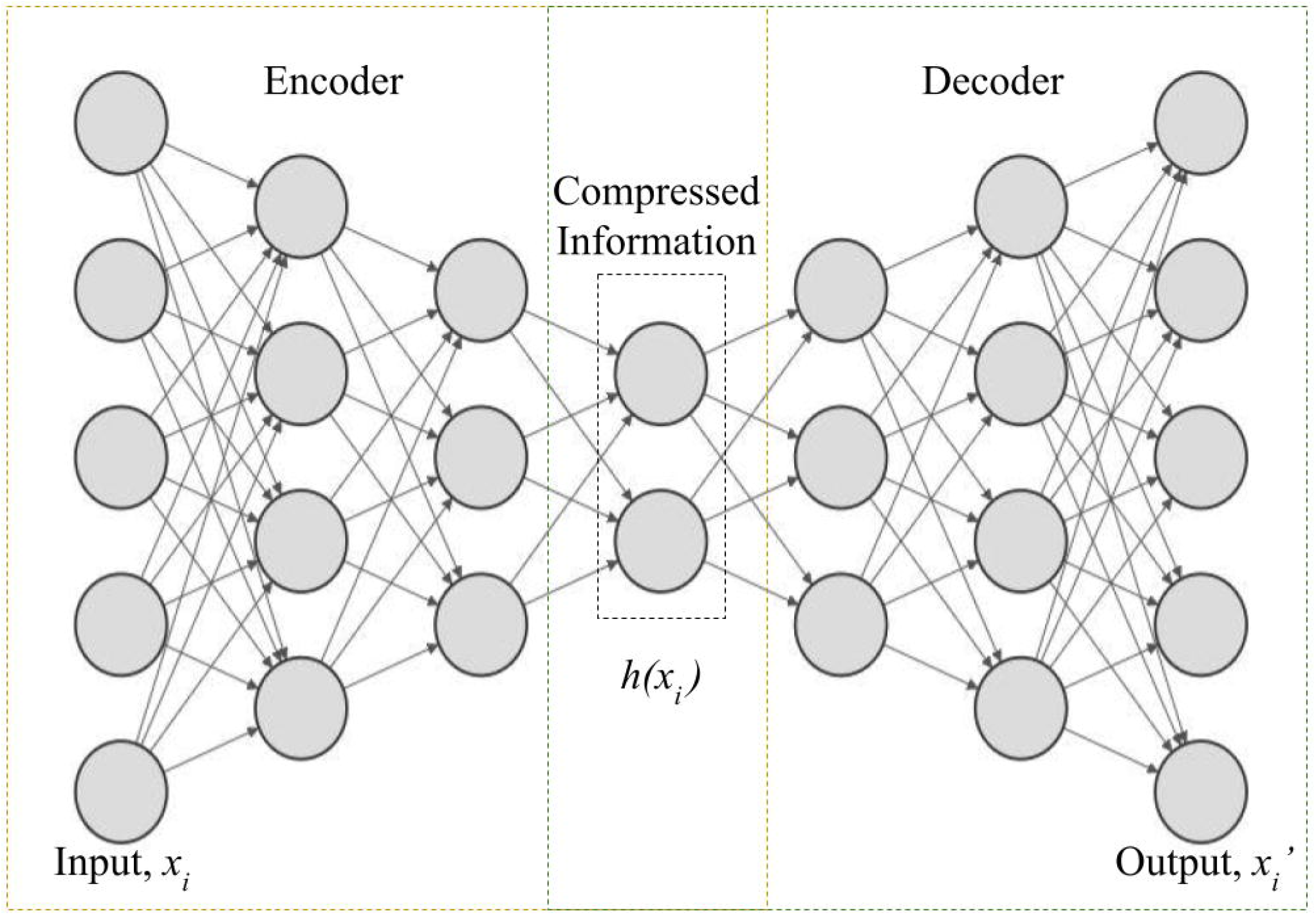
Architecture of a Deep Autoencoder.

The activation function used for each layer, except the middle and decoder layers, is “ReLU” (Patterson and Gibson, 2017), which scales negative output value to zero:

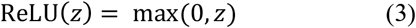

For the middle and decoder layers, the activation function is a “sigmoid” (Patterson and Gibson, 2017), which scales the output to the range [0, 1]:

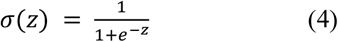

The reconstruction error was calculated as mean squared error (MSE) function:

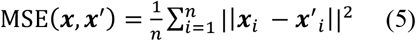

where ***x***_*i*_ and ***x***′_*i*_ are the measured and predicted values, respectively, and *n* is the number of samples with *i* ∈ [1, *n*].

An autoencoder model was used to compress the genome-wide polymorphism data. Each dataset was split into training (60%), testing (20%), and validation (20%) sets using ‘train_test_split’ function of scikit-learn. To optimize the performance of compression model, we employed a ‘KerasRegressor’ wrapper from Keras to tune hyperparameters (Table 1) via scikit-learn’s RandomizedSearchCV.’ For the C7AIR dataset, hyperparameter tuning was performed on the entire dataset because of its smaller dimensionality. In contrast, for the HDRA dataset, tuning was performed on a small subset of the training data, specifically the first 1000 splits of data, each containing 1568 × 28 samples.

**Table 1.**
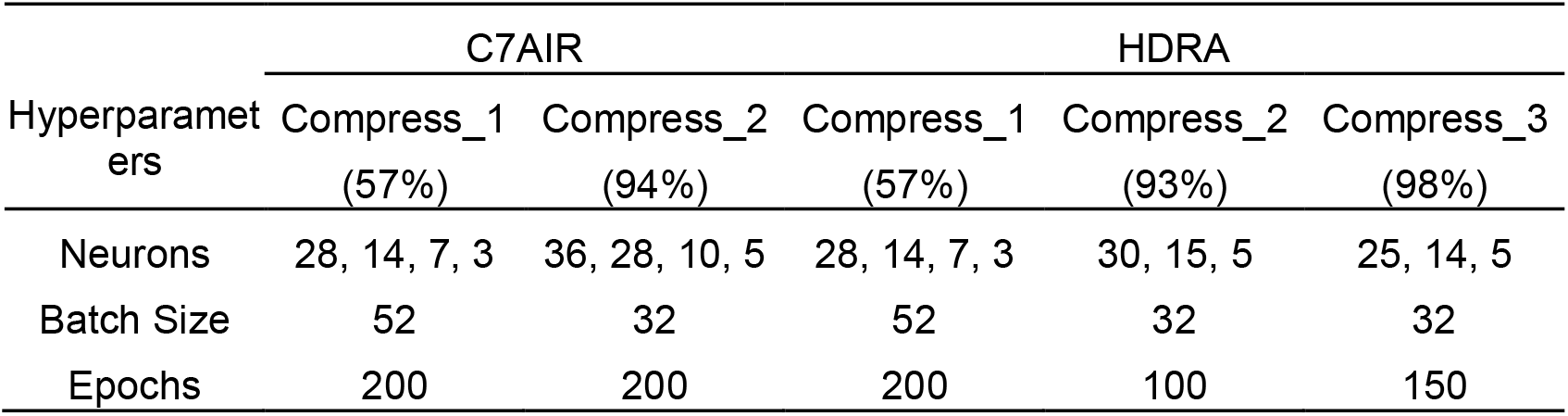
Hyperparameters determined for the C7AIR and HDRA datasets. The learning rate, batch size, and loss function hyperparameters were tuned by exploring the values slightly above and below TensorFlow’s default range. The optimum learning rate for the Adam optimizer was selected from the logarithmic scale of commonly used values: {0.01, 0.001, and 0.0001}. Batch sizes of {16, 32, 52, and 64} were tested, and the optimal loss was determined by comparing three different loss functions: mean squared error, binary cross-entropy, and mean absolute error.

Although the architecture of the autoencoders was reported in our previous work (Islam et al., 2023), it is described here for the sake of completeness. For C7AIR genotype data, the selected model consisted of three hidden layers in both the encoder and decoder networks. In Compress_1, the input layer of the network had 28 nodes, followed by 14 and seven nodes in the hidden layers, with a code size of 3. The output from Compress_1 was used as the input for Compress_2, which had an input layer of 36 nodes, hidden layers of 28 and 10 nodes, and a code size of five. Both compressions were trained using the Adam optimizer with a learning rate of 0.001. ReLU activation was applied to all layers, except the middle and last layers, where sigmoid activation was used. The model was trained with the MSE loss using a mini-batch size of 52 for Compress_1 and 32 for Compress_2 with 200 epochs for both.

For the HDRA genotype data, the architecture was similar, but adapted to the higher dimensionality of the dataset. Compress_1 has layers with [28, 14, 7, 3] nodes; Compress_2 had [30, 15, 5] nodes; and Compress_3 had [25, 14, 5] nodes. Compress_1 was trained with 200 epochs and a batch size of 52, Compress_2 was trained with 100 epochs and a batch size of 32, and Compress_3 was trained with 150 epochs and a batch size of 32. Other parameters were identical to those used for the C7AIR data.

The compression model was implemented using the Keras functional API [41] built on TensorFlow in Python. For the C7AIR data, the compression levels were Compress_1 (57%) and Compress_2 (94%), indicating that Compress_1 reduced the data size to 43% of the original data, whereas Compress_2 reduced it to 6%. Similarly, for the HDRA data, Compress_1 (57%), Compress_2 (93%), and Compress_3 (98%) were applied, reducing the data size to 43%, 7%, and 2% of the original size, respectively. The mean squared error (MSE) loss for the C7AIR genotype data was 0.051 for Compress_1 and 0.039 for Compress_2. Similarly, for the HDRA genotype data, the MSE losses remained low, with values of 0.0156, 0.0749, and 0.0911 for Compress_1, Compress_2, and Compress_3, respectively, indicating that high-quality compression was achieved with minimal loss.

### 2.3 Prediction Model using CNN

The CNN architecture (Figure 2) was designed to handle input variables distributed in spatial patterns, such as one-dimensional (e.g., SNPs or text), two-, or three-dimensional (e.g., images). CNNs are neural networks that learn higher-order features using convolution operations instead of full-matrix multiplications in hidden layers (Goodfellow et al., 2016). A typical CNN comprises both dense and fully connected layers, as well as convolutional layers. In each convolutional layer, a convolution operation was applied along the input using predefined widths and strides. These convolutional processes, known as “kernels” or “filters,” function similarly to neurons in an MLP. The output of the convolution is expressed as an integral transformation (Sandhu et al., 2020) as follows:

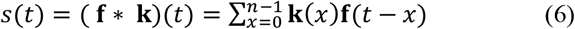

Here, **k** represents the kernel and the convolution describes the transformation of **f** into *s*(*t*). This operation shifts **f** over the kernel along each chromosome, allowing the filters to account for the linkage disequilibrium across the chromosome.

**Figure 2.**
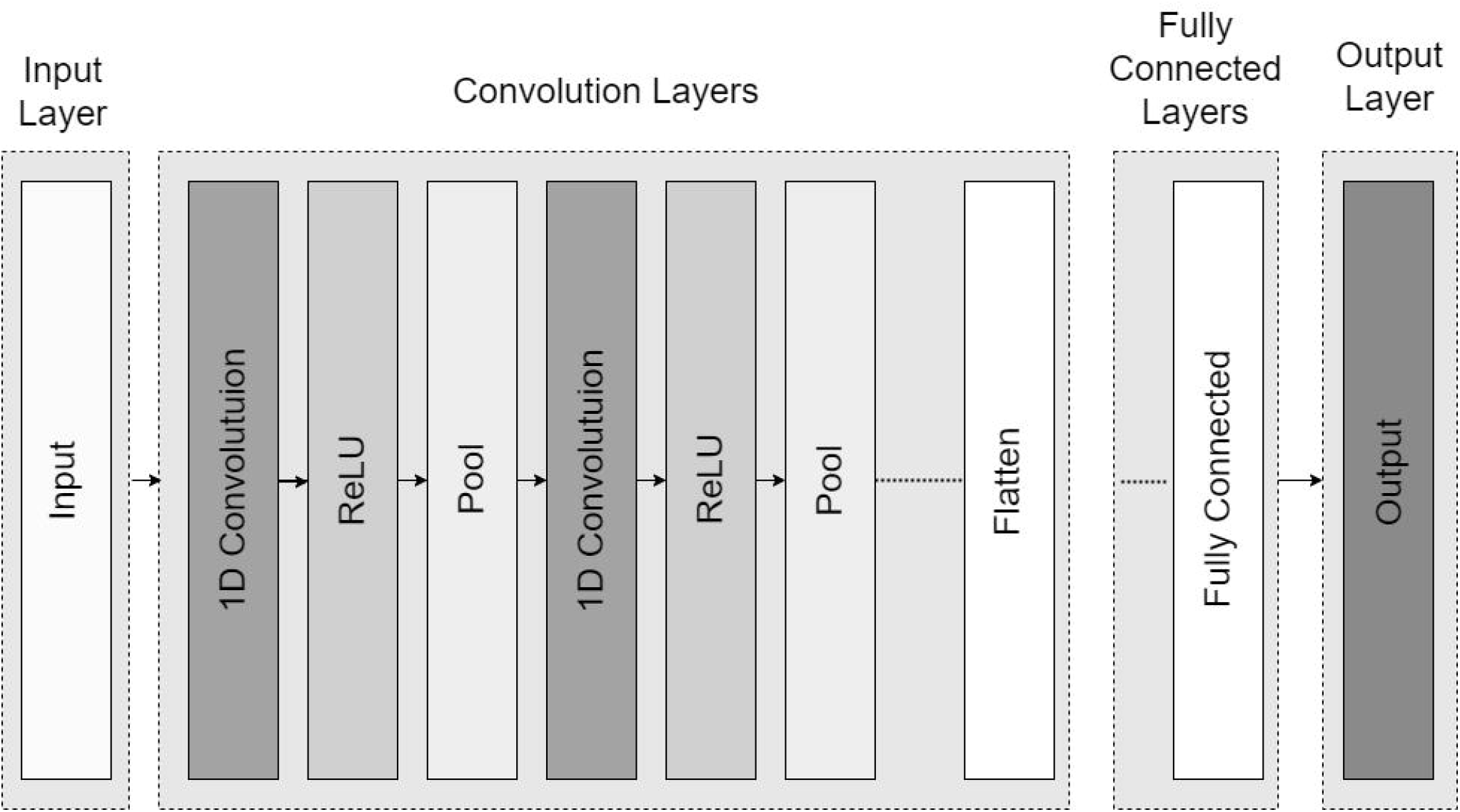
Basic Architecture of a 1D Convolutional Neural Network (CNN).

To generate the output, an activation function was applied after each convolution. A “pooling” operation is typically used to smooth the result by taking the mean, maximum, or minimum of the kernel outputs at successive positions and merging them. One key advantage of convolutional networks is their ability to reduce the number of parameters to be estimated, while also featuring sparse interactions and translation equivariance. CNNs comprise three main types of layers: convolutional, pooling, and fully-connected (FC) layers (Goodfellow et al., 2016). The CNN architecture was constructed by stacking the layers. Additionally, two important components, the dropout layer and activation function, play a critical role in defining the performance of the model.

The proposed CNN architecture was designed to process 1D input data for regression tasks. It begins with four convolutional layers, with filter sizes of 32, 64, 96, and 256, and kernel sizes of 3, 7, 5, and 5. Each convolutional layer is followed by a LeakyReLU activation function and batch normalization to improve stability and accelerate convergence. MaxPooling1D layers with a pool size of 2 were applied after each convolutional layer to reduce the spatial dimensions and prevent overfitting. Dropout layers with rates of 0.4, 0.4, 0.3, and 0.3, were placed after each BatchNormalization layer to further mitigate overfitting by randomly dropping neurons during training. The model was then transitioned to two fully connected layers with 256 and 224 units, respectively, followed by LeakyReLU activation and batch normalization. The final output layer is a dense layer with a single unit that provides a regression prediction. This architecture integrates convolutional, activation, normalization, pooling, and dropout layers to efficiently process and predict 1D data inputs.

For genetic value prediction using CNN, we used the same two datasets, C7AIR and HDRA, as in our deep-learning compression-based prediction model. The CNN was applied to both the original uncompressed data and various levels of compressed data. The estimated genotypic values of a target trait with omitted missing entries were arranged in the same order as that of the original and compressed datasets.

We divided the datasets into two subsets using the ‘train_test_split’ function of scikit-learn: 80% for training and 20% for validation. The model was trained on the training subset, and its performance was evaluated at regular intervals on the validation subset. In the following section, we outline the process of determining the optimal parameters for training a CNN model.

### 2.4 Determination of Hyper-parameters

Hyperparameter tuning is crucial for optimizing CNN models given the wide range of possible parameter combinations. In this study, we used a hyperband tuner for efficient hyperparameter optimization with the aim of minimizing the MSE across all evaluated traits. The model was implemented using Keras and incorporated layers such as Conv1D, LeakyReLU, BatchNormalization, MaxPooling1D, Flatten, Dense, and Dropout to reduce overfitting. Hyperparameter optimization was performed using a hyperband tuner, which iteratively adjusted the key parameters, including the number of filters, kernel sizes, dropout rates, number of convolutional layers, dense layer units, and learning rates. The search space included filters (32 to 256), kernel sizes (3, 5, 7), dropout rates (0.3 to 0.7), dense layer units (256 to 512 and 128 to 256), and learning rates (1e-4 to 1e-2). The data were divided into training and validation sets, with callbacks, such as EarlyStopping and ReduceLROnPlateau, used to prevent overfitting and ensure efficient training. Optimal hyperparameters, such as specific filter sizes, kernel sizes, dropout rates, and learning rates, were selected based on the lowest validation loss and were used to build the final CNN model for genetic value prediction. The selected hyperparameters are listed in Table 2.

**Table 2.**
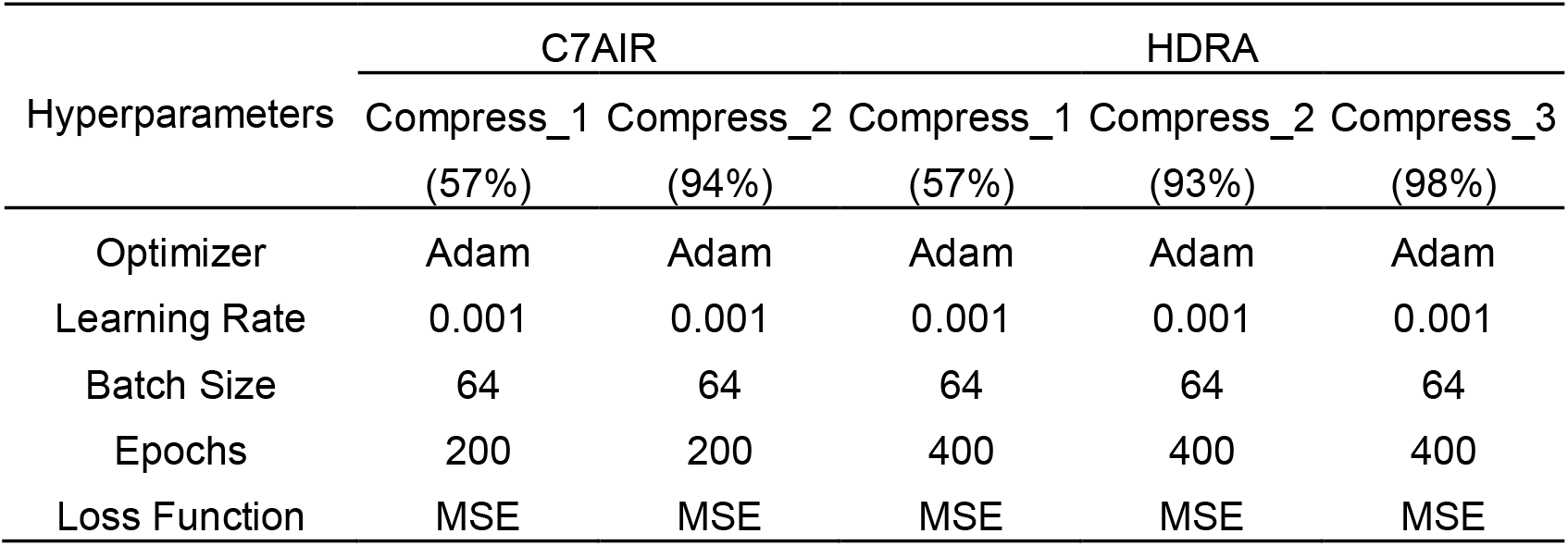
Hyperparameters determined for the C7AIR and HDRA datasets using a RandomizedSearchCV for the CNN model.

| Hyperparameters | C7AIR |  | HDRA |  |  |
| --- | --- | --- | --- | --- | --- |
|  | Compress_1 | Compress_2 | Compress_1 | Compress_2 | Compress_3 |
|  | (57%) | (94%) | (57%) | (93%) | (98%) |
| Optimizer | Adam | Adam | Adam | Adam | Adam |
| Learning Rate | 0.001 | 0.001 | 0.001 | 0.001 | 0.001 |
| Batch Size | 64 | 64 | 64 | 64 | 64 |
| Epochs | 200 | 200 | 400 | 400 | 400 |
| Loss Function | MSE | MSE | MSE | MSE | MSE |

## 3 Results

### 3.1 Prediction of genotypic values based on the compressed data using CNN

To evaluate the predictive accuracy of the CNN model and address the impact of data compression, we applied the CNN models to the compressed datasets. Figure 3 illustrates the prediction accuracy of the CNN across various compression levels for the C7AIR and HDRA datasets. Figure 3A shows the performance of the CNN for plant height in the C7AIR dataset at three different compression levels. The model achieved an accuracy rate of 80% at 0% compression (uncompressed data). As the compression increased to 57%, the accuracy decreased slightly to 75%, and further dropped to 74% at 94% compression. Despite the data reduction, the CNN maintained a relatively stable and high level of accuracy, demonstrating its robustness in handling compressed genomic data for prediction.

**Figure 3.**
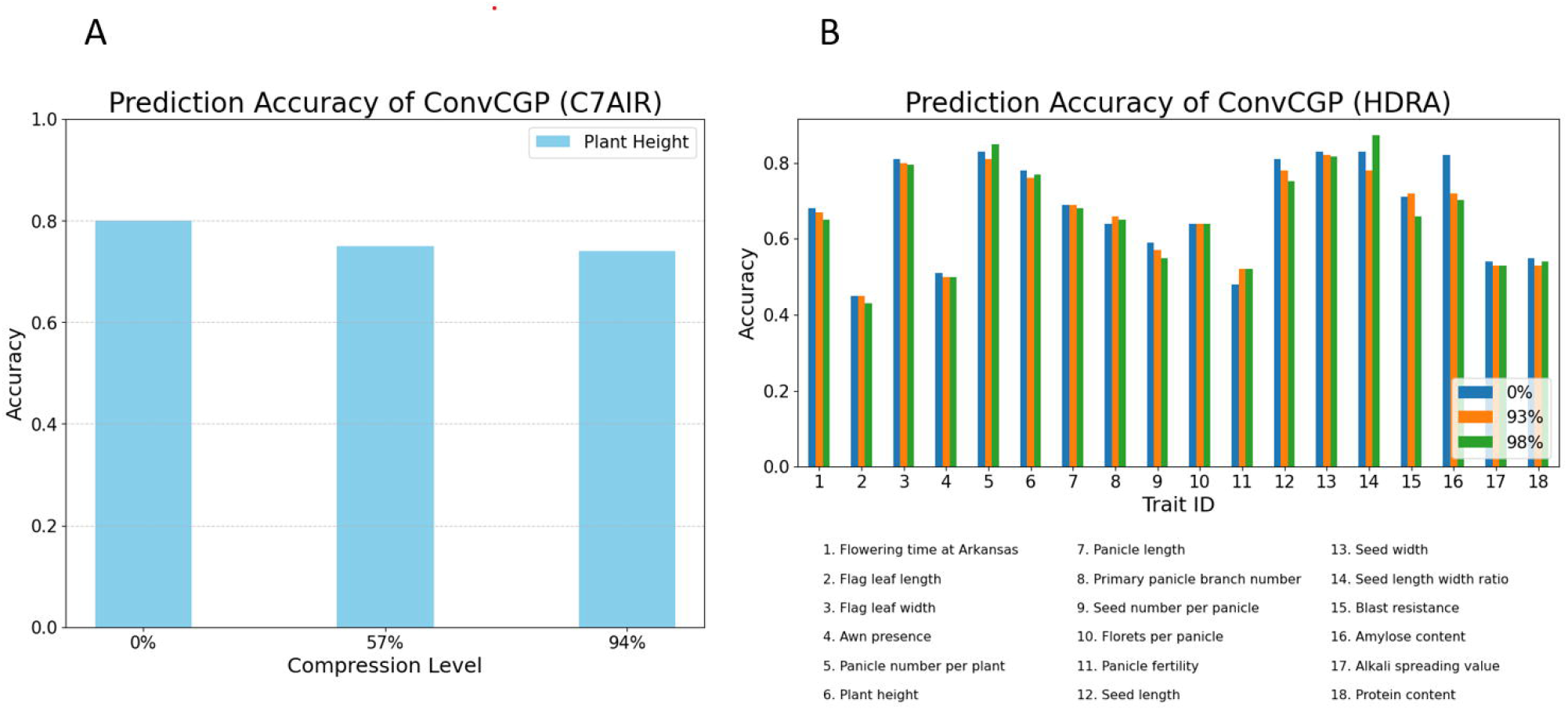
ConvCGP prediction accuracy for (A) C7AIR and (B) HDRA dataset.

Figure 3B shows the HDRA dataset, highlighting traits that maintained a high prediction accuracy, even after substantial compression. Notably, traits such as flag leaf width (trait 3), panicle number per plant (trait 5), seed width (trait 13), and blast resistance (trait 15) consistently exhibited strong performances across all compression levels. These traits exhibit minimal changes in accuracy compared with predictions from uncompressed data, demonstrating the ability of the CNN to preserve prediction quality despite a substantial reduction in data dimensions.

Other traits, including plant height (trait 6), seed number per panicle (trait 9), and seed length (trait 12), also maintain stable accuracy, demonstrating the effectiveness of the CNN in preserving predictive performance across various traits. Even at high compression levels, the CNN model outperformed the other prediction methods, reaffirming its robustness and adaptability for handling compressed genomic data across a wide range of traits. This suggests that CNNs are highly suitable for predictive modeling in scenarios where data storage or computational resources are limited, because they can deliver high-quality predictions from significantly compressed data.

The prediction times for the ConvCGP method across the C7AIR and HDRA datasets were significantly reduced when compressed data were used. For the C7AIR dataset, the prediction time significantly decreased as the compression level increased. At 0% compression, ConvCGP required 2 min and 50 s, whereas at 57% compression, the time decreased to 53 s. When the data were further compressed to 94%, the prediction time reduced to 14 s. This trend was consistent in the HDRA dataset. Starting at 0% compression, ConvCGP required 1 d, 1 h, and 6 min on average per trait prediction. However, with 93% compression, this time decreases to 54 min and 45 s, and further compression to 98% results in prediction times of 39 min and 10 s, respectively. These results illustrate the effectiveness of data compression in improving computational efficiency. Even at high compression levels, the quality of predictions remained intact, as indicated by the high levels of information retained across both datasets. Thus, applying data compression before prediction not only accelerates the processing time but also ensures that essential information is preserved, facilitating faster and equally reliable predictions.

All experiments in this study were conducted on a PC with an Intel(R) Core (TM) i9-10980XE, 3.00 GHz CPU, 128 GB RAM, GPU RTX 3090, and 64-bit Windows 10 Pro operating system.

### 3.2 Comparing prediction performance with other methods

We conducted a comparative analysis of ConvCGP against two recent methodologies that rely on kinship matrices and principal component (PCs) scores, as well as our previous framework, DeepCGP. Kinship matrices (Nazzicari and Biscarini, 2022), such as the realized additive (*K*_A_), dominance (*K*_D_), additive-by-additive (K_AA_), additive-by-dominance (K_AD_), and dominance-by-dominance (*K*_*DD*_), represent genetic relationships. SNPs are filtered based on minor allele frequency (MAF), and the five genetic relationships (K_A_, K_D_, K_AA_, K_AD_, K_DD_) yield 15 kinship matrices. These kinship matrices condensed representations of genetic relatedness derived from genome-wide marker genotype data, reduce input dimensionality and, when stacked within a 2D-CNN, leverage genetic similarities to improve predictions for continuous genotypic values of traits. In our datasets, we only used K_A_ and K_AA_ matrices, as the HDRA dataset contains few heterozygous genotypes. Since the dominance effect primarily affects heterozygous genotypes, its inclusion had minimal impact on the overall results. Another approach using PCs involves principal component analysis to reduce input dimensions, followed by multimodal deep learning (DL) techniques. These techniques have shown comparable prediction accuracy between DL and GBLUP (Kick et al., 2023). The PCs captured 31% of the variance with 8 PCs, 50% with 50 PCs, and over 99% with 1,725 PCs. For the PCs analysis, we removed one negative eigenvalue from the HDRA data and reduced the dimensionality. In addition to these kinship matrices and PC-based approaches, we compared our method with our previous work, DeepCGP, a compression-driven predictive model that uses deep learning autoencoders for data compression and random forest regression for prediction (Islam et al., 2023).

Figure 4 illustrates the prediction accuracy of the four models --kinship, PCs, DeepCGP, and ConvCGP--across 18 traits using a 98% compressed version of the HDRA dataset. The vertical axis represents prediction accuracy, whereas the horizontal axis lists the traits. ConvCGP consistently outperformed the other models for most traits, demonstrating its ability to effectively handle highly compressed data. Notably, for traits such as flowering time in arkansas (trait 1), panicle number per plant (trait 5), plant height (trait 6), and seed length-width ratio (trait 14), ConvCGP achieved significantly higher accuracy than Kinship, PCS, and DeepCGP. ConvCGP also excels in predicting key quality traits, such as blast resistance (trait 15) and amylose content (trait 16), where it approaches near-perfect accuracy. These results indicate that the ConvCGP is particularly skilled at capturing intricate genetic patterns, even when the dataset is compressed to retain only 2% of the original information. Whereas DeepCGP performed well overall, ConvCGP surpassed it for most traits, showcasing the additional advantages of convolutional layers in extracting meaningful patterns from reduced-dimensional data. In contrast, Kinship and PCs exhibited lower accuracy, particularly for traits such as blast resistance (trait 15) and seed composition (traits 9, 12, 13, and 14), highlighting their limitations in modeling complex genetic relationships. Despite its high level of compression, the consistent performance of the ConvCGP across all traits highlights its robustness and adaptability, making it a powerful tool for predicting a wide range of genetic traits with minimal data. This analysis strongly supports the superiority of ConvCGP over conventional methods even in situations where extreme data compression is necessary.

**Figure 4.**
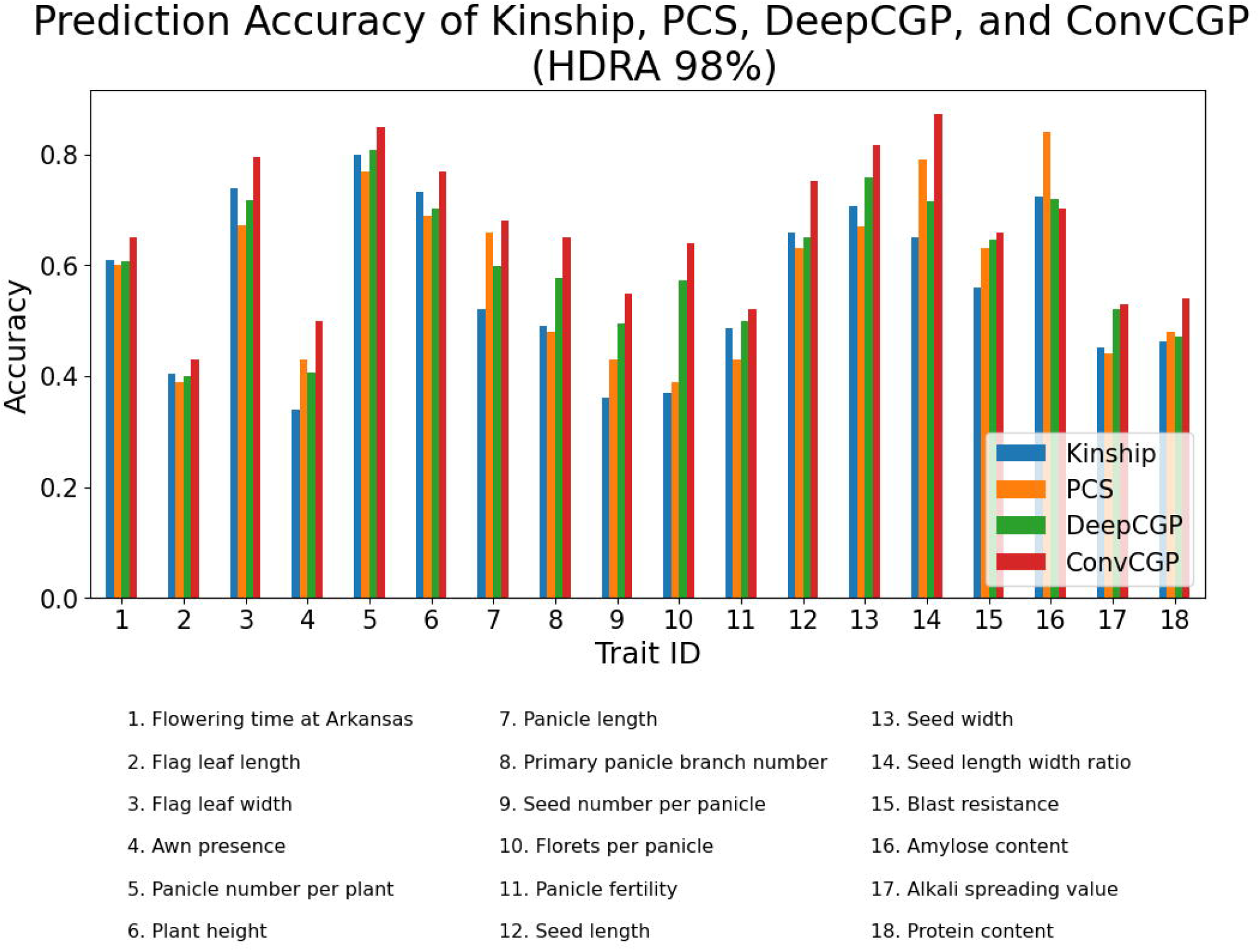
Prediction accuracy of Kinship, PCs, DeepCGP and ConvCGP for the HDRA dataset.

## 4 Discussion

Efficient use of high-dimensional genome-wide polymorphism data for plant and animal breeding requires innovative platforms, which significantly reduce the resources required for storage (Islam et al., 2023) and processing. Few studies have focused on reducing the dimensions of genome-wide polymorphism data while simultaneously predicting genotypic values for a target trait. The kinship matrix, PC, and DeepCGP approaches were examined. Although both kinship matrices and PCs successfully reduced dimensionality and enabled predictions from compressed data, their prediction accuracy remained low. Conversely, in our previous study, DeepCGP, which combined deep learning for compression with machine learning for prediction, outperformed kinship matrices and PC-based methods. However, we identified the potential to further enhance prediction accuracy by adopting a DL-based prediction method rather than relying on machine learning for the prediction phase, as in DeepCGP.

In this study, we proposed ConvCGP, a deep-learning-based method that improves prediction accuracy by utilizing CNNs to predict compressed data. This approach was applied to rice by employing CNN architectures with convolutional, pooling, and fully connected layers to predict the genotypic values of traits from compressed genome-wide polymorphism data. The convolutional layers, consisting of multiple convolution kernels, generated various feature maps from the input representations. Each neuron in the feature map was connected to a specific region in the neighboring neurons of the previous layer.

The CNN architecture was constructed by stacking layers. During training, we optimized the algorithm and selected the network parameters. With a relatively shallow network (three or five layers) and carefully tuned parameters, we achieved 99.3% relative accuracy using genome-wide polymorphism data that had been compressed by 98%, reducing the data size to only 2% of its original size. This level of accuracy was comparable to that obtained using the original data, demonstrating the effectiveness of ConvCGP in predicting genotypic values from highly compressed genome-wide polymorphism data.

We developed a compression-based genomic prediction model, ConvCGP, using deep learning, which enhanced breeding efficiency while significantly reducing the storage requirements for genome-wide polymorphism data. The key advantage of using DL for compression is its ability to extract meaningful information from the genetic architecture, effectively modeling complex patterns with fewer computational resources compared to conventional approaches. The experimental results are promising, achieving high accuracy of genotypic value prediction while maintaining robustness, even with compressed data. Our model allows flexibility in the compression levels, which can be adjusted based on storage constraints, the time required to construct a CNN, or accuracy requirements. ConvCGP outperformed kinship-matrix-based approaches and DeepCGP in terms of predictive performance. However, this approach has only been tested on rice germplasm data, and its generalizability to other datasets, including those involving animals or other plant species, has not yet been evaluated. Therefore, researchers should apply the ConvCGP with caution when using it for other datasets. Future work will include analyzing the gradients within the neural network that predict genotypic values from high-dimensional genome-wide polymorphism data, and exploring the potential of deep learning-based compression methods to identify important SNP sets.

## 5 Conclusions

In summary, ConvCGP, a pioneering DL model, was introduced as an innovative solution for compressing high-dimensional genome-wide polymorphism data while accurately predicting genotypic values from the compressed information. ConvCGP demonstrates great potential for complex modeling, particularly through the novel application of deep learning in genomic prediction. Using compressed data as input variables, this approach significantly improves the computational efficiency of CNN-based deep learning. In addition, ConvCGP provides a robust solution for handling high-dimensional genomic data, maintaining high prediction accuracy even with compressed inputs. It also offers advantages in terms of storage efficiency and reduced computational time, making it a promising tool for genomic prediction and selection in both plant and animal breeding.

## 8 Author Contributions

H. Shimono initiated the project. T. Islam and C. H. Kim designed the study. T. Islam conducted all the analyses and wrote the main manuscript with the support of H. Iwata. C. H. Kim, A. Kimura, and H. Iwata supervised the study. All the authors reviewed the manuscript.

## 10 Data Availability Statement

Our code and data are available at https://github.com/tanzilamohita/ConvCGP.

